# CMKLR1-targeting peptide tracers for PET/MR imaging of breast cancer

**DOI:** 10.1101/575902

**Authors:** Sarah Erdmann, Lars Niederstadt, Eva Jolanthe Koziolek, Juan Daniel Castillo Gómez, Sonal Prasad, Asja Wagener, Jan Lennart von Hacht, Sandy Hallmann, Samantha Exner, Sebastian Bandholtz, Nicola Beindorff, Winfried Brenner, Carsten Grötzinger

## Abstract

Molecular targeting remains to be a promising approach in cancer medicine. Knowledge about molecular properties such as overexpression of G protein-coupled receptors (GPCRs) is thereby offering a powerful tool for tumor-selective imaging and treatment of cancer cells. We utilized chemerin-based peptides for CMKLR1 receptor targeting in a breast cancer xenograft model. By conjugation with radiolabeled chelator 1,4,7,10-tetraazacyclododecane-1,4,7,10-tetraacetic acid (DOTA), we obtained a family of highly specific and affine tracers for hybrid *in vivo* imaging with positron emission tomography (PET)/ magnetic resonance (MR) and concomitant biodistribution studies.

**Methods:** We developed five highly specific and affine peptide tracers targeting CMKLR1 by linker-based conjugation of chemerin peptide analogs (CG34 and CG36) with radiolabeled (^68^Ga) chelator DOTA. Our established xenograft model with target-positive DU4475 and negative A549 tumors in immunodeficient nude mice enabled CMKLR1-specific imaging *in vivo*. Therefore, we acquired small animal PET/MR images, assessed biodistribution by *ex vivo* measurements and investigated the tracer specificity by blocking experiments.

**Results:** The family of five CMKLR1-targeting peptide tracers demonstrated high biological activity and affinity *in vitro* with EC_50_ and IC^50^ values being below 2 nM. Our target-positive (DU4475) and target-negative (A549) xenograft model could be confirmed by *ex vivo* analysis of CMKLR1 expression and binding. After preliminary PET imaging, the three most promising tracers ^68^Ga-DOTA-AHX-CG34, ^68^Ga-DOTA-KCap-CG34 and ^68^Ga-DOTA-ADX-CG34 with apparent DU4475 tumor uptake were further analyzed. Hybrid PET/MR imaging along with concomitant biodistribution studies revealed distinct CMKLR1-specific uptake (5.1% IA/g, 4.5% IA/g and 6.2% IA/g 1 h post-injection) of our targeted tracers in DU4475 tumor tissue. More strikingly, the tumor uptake could be blocked by excess of unlabeled peptide (6.4-fold, 7.2-fold and 3.4-fold 1 h post-injection) and further confirmed the CMKLR1 specificity. As our five tracers, each with particular degree of hydrophobicity, showed different results regarding tumor uptake and organ distribution, we identified these three tracers with moderate, balanced properties to be the most potent in receptor-mediated tumor targeting.

**Conclusion:** With the breast cancer cell line DU4475, we established a model endogenously expressing our target CMKLR1 to evaluate our chemerin-based peptide tracers as highly affine and specific targeting agents. Eventually, we demonstrated the applicability of our ^68^Ga-labeled tracers by visualizing CMKLR1-positive breast cancer xenografts in PET/MR imaging and thus developed promising theranostics for tumor treatment.

## Introduction

Molecular targeting remains to be one of the most promising approaches in cancer diagnosis and therapy. Targeted tracers in combination with highly sensitive positron emission tomography (PET) may facilitate early tumor recognition and staging staging as well as therapeutic stratification and response monitoring. While the most widely used PET tracer ^18^F-fluoro-deoxyglucose (^18^F-FDG) enables tumor detection by glucose metabolic imaging and hence also visualizes non-neoplastic cells and tissues, tumor-specific targeting is aiming to provide more sensitive and precise imaging results with less non-specific background [1]. Preferred molecular targets are considered to localize at the cell surface or within the extracellular matrix (ECM), as tracers can reach them without crossing the cell membrane. Consequently, transmembrane receptors, transporters and other antigens associated with the cell membrane as well as ECM proteins such as fibronectin or tenascin can be regarded as potential targets [2]. With either high overexpression or even exclusive target expression in tumor cells, appropriately targeted tracers may lead to improved tumor-to-background ratios and thus higher diagnostic sensitivity and specificity.

Several G protein-coupled receptors (GPCRs) have been identified as tumor-specific targets. Beside other well-characterized cell surface proteins such as growth factor receptors (e.g. epithelial growth factor receptor, EGFR [3]), transmembrane glycoproteins (e.g. prostate-specific membrane antigen, PSMA [4]) and transmembrane receptors (e.g. integrins [5]), GPCRs have emerged as molecular targets in the field of cancer care [6, 7]. Thus, upregulated receptor expression in cancer cells can be utilized for tracer-based tumor imaging, for pharmacological intervention through GPCR signaling modification and for peptide receptor radionuclide therapy (PRRT). Unlike antibodies, small-sized receptor ligands such as peptides have a number of advantages regarding their *in vivo* circulation, biodistribution, tumor penetration and low immunogenic potential [8-11].

One of the earliest successful receptor-based strategies in cancer targeting was the use of somatostatin analogs (SSA) for diagnostic imaging and treatment of somatostatin receptor-bearing tumors. For human somatostatin receptor (SSTR), a G protein-coupled transmembrane glycoprotein, five subtypes (SSTR1-SSTR5) are known to be expressed in neuroendocrine tumors (NETs) [12, 13]. With a prevalence of 80 to 100% and the highest abundance for SSTR2 expression, NETs of the gastroenteropancreatic tract (75%, compared to 25% NETs in lungs) are suited for SSTR targeting [14, 15]. While ^111^In-pentetreotide (^111^In-DTPA-octreotide) for single photon emission computed tomography (SPECT) was the first SSTR tracer with widespread clinical use [16, 17], ^68^Ga-DOTATOC (^68^Ga-DOTA-Tyr^3^-octreotide) for PET and ^177^Lu-DOTATATE (^177^Lu-DOTA-Tyr^3^-octreotate) for PRRT have now become available [18-20].

Chemokine-like receptor 1 (CMKLR1), a G protein-coupled receptor, and its ligand chemerin are known to be involved in inflammation, adipogenesis and glucose metabolism [21, 22]. Dysregulation of the receptor-ligand system has been linked to several pathologies such as metabolic syndrome and cardiovascular diseases [23, 24]. Apart from a study demonstrating recruitment of immune cells and hence an antitumor response in melanoma, the role of the chemoattractant chemerin and its receptors in cancer is largely unknown [25, 26]. Our lab has recently characterized CMKLR1 as a target overexpressed in breast cancer, esophageal squamous cell carcinoma (ESCC) and pancreatic adenocarcinoma (manuscript in preparation). Increased CMKLR1 expression, chemerin-mediated tumor cell migration and invasion in ESCC were also found by Kumar et al. [27]. Likewise, CMKLR1 has been implicated in neuroblastoma proliferation [28], hepatocellular carcinoma metastasis [29] and migration/invasion of gastric cancer [30]. CMKLR1 may therefore be a promising target for molecular imaging and targeted radionuclide therapy in overexpressing tumor entities. This study evaluated the potential of a family of novel CMKLR1 peptide-DOTA tracers for PET/MR imaging in a breast cancer animal model.

## Material and Methods

### Peptides

All peptides and DOTA-peptide conjugates were from peptides&elephants (Hennigsdorf, Germany). They were analyzed by mass spectrometry to confirm the presence of the correct molecular mass. Peptides and peptide conjugates were used at a purity of greater than 95%. Analysis data for peptides and peptide conjugates have been deposited in an open data repository for public access: http://doi.org/10.5281/zenodo.2591417

### Cell culture

For *in vitro* analysis, HEK293a stably transfected with huCMKLR1 and G^α16^ were used. For the *in vivo* mouse model, the human breast cancer cell line DU4475 with endogenous CMKLR1 expression and the target-negative human lung carcinoma cell line A549 were used. All cell lines were obtained from ATCC/LGC Standards (Wesel, Germany) and were cultured in RPMI1640 medium containing 10% fetal bovine serum (both from Biochrom, Berlin, Germany) and, in case of transfected cells, selection agents G418 (Biochrom) and zeocin (Invitrogen, Carlsbad, USA).

### Xenografts

For *in vivo* experiments, at least 8-week-old female athymic NMRI*-Foxn1*^*nu*^ */Foxn1*^*nu*^ mice (Janvier Labs, Saint-Berthevin, France) were used. Animal care followed institutional guidelines and all experiments were approved by local animal research authorities. For the generation of tumor xenografts, 5 × 10^6^ cells of both DU4475 and A549 cells were inoculated subcutaneously into the left and right shoulder, respectively (1:1 PBS/Matrigel Basement Membrane Matrix High Concentration, Corning, Corning, USA). Tumors were allowed to grow for two to four weeks (tumor volume > 100 mm^3^) after cell inoculation.

### Immunofluorescence

For immunofluorescent staining of tumor tissue, the frozen xenografts were embedded in Tissue-Tek O.C.T Compound (Sakura Finetek, Torrance, USA), trimmed by cryostat (12-18 µm) and transferred on glass slides (Superfrost, Thermo Fisher Scientific, Waltham, USA). Xenograft sections were fixed with 1:1 methanol/acetone for two minutes and air-dried. After washing with PBS (Biochrom, Berlin, Germany), sections were blocked with 2% milk powder (blotting grade; Bio-Rad Laboratories, Hercules, USA) in PBS for 30 minutes. Sections were incubated with the primary anti-CMKLR1 antibody developed in our lab (#21-86; polyclonal from rabbit, immunogenic peptide sequence: SSWPTHSQMDPVGY; 3 µg/mL diluted in 0.1% BSA in PBS) in a wet chamber over night at 4 °C. After washing, sections were incubated with the secondary antibody goat-anti-rabbit-Cy2 (Jackson ImmunoResearch, West Grove, USA; 7.5 µg/mL diluted in 0.1% BSA in PBS) for one hour. After washing with TBS, nuclei staining with 1 µM SytoxOrange (Thermo Fisher Scientific, Waltham, USA) in TBS followed. Finally, the cryosections were fixed with 96% ethanol for two minutes, embedded with Immu-Mount (Thermo Fisher Scientific, Waltham, USA) and analyzed with a confocal laser-scanning microscope (LSM510, Carl Zeiss, Jena, Germany).

### Calcium mobilization assay

Optical ViewPlate 96-well microplates (PerkinElmer, Waltham, USA) were coated with poly-D-lysine (10 µg/mL, 50 µL/well) for 30 minutes at 37 °C, washed with PBS and air dried before seeding 40,000 HEK293a huCMKLR1 G_α16_ cells per well. The next day, cells were starved with serum-free medium for 30 minutes and incubated with loading medium consisting of serum-free medium with 2.5 mM probenicid (Sigma-Aldrich, St. Louis, USA) and 2 µM Fluo-4 AM (Thermo Fisher Scientific, Waltham, USA). After a loading time of 45 minutes, cells were washed with 100 μL C1 buffer (130 mM NaCl, 5 mM KCl, 10 mM Na-Hepes, 2 mM CaCl^2^, 10 mM glucose) followed by 20 minutes of incubation in the dark. After another washing and incubation cycle, the cell plate was placed into the CellLux Calcium Imaging System (PerkinElmer, Waltham, USA) and the assay was performed automatically according to the following protocol. After a baseline measurement for 30 seconds, the ligands (prepared in C1 buffer with 0.5% BSA in double concentration and pipetted in a U-bottom plate) were added on top of the cells and the fluorescence intensity was recorded for a further 60 seconds. Obtained raw data were analyzed using AssayPro (PerkinElmer, Waltham, USA). After spatial uniformity correction, maximum response values (F) were normalized to baseline values (F^0^) of each well using the following equation: response = ΔF/F_0_ = (F – F_0_)/F_0_.

### Iodination of chemerin-9

Radioactive iodination of chemerin-9 was performed by the chloramine T method [31]. For labeling, 10 nmol of chemerin-9 in 25 μL iodination buffer (0.5 M sodium phosphate, pH 7.4) were mixed with 1 mCi carrier-free Na^125^I (NEZ033L010MC, PerkinElmer, Waltham, USA) in an HPLC glass vial with microvolume insert. The reaction was started by adding 4 μL chloramine T (1 mg/mL in water). After 20-30 seconds, 4 μL of sodium metabisulfite (2 mg/mL in water) were added to stop the iodination. HPLC purification was performed to separate unlabeled from labeled radioactive peptide on an Agilent ZORBAX 300 Extend-C18 column using a gradient from 20 to 50% acetonitrile (+0.1% TFA) against water (+0.1% TFA) for 20 minutes. First, 1-2 μL of the reaction mixture were preanalyzed to determine the retention time of the radioactive peptide. This fraction was then collected during the main run, diluted with radioactive binding buffer to prevent radiolysis, aliquoted and stored at −20 °C.

### Competitive binding studies

Competitive binding studies were performed with cell membrane preparations. Therefore, cellular monolayer cultures were washed with pre-warmed PBS and dissociated in ice-cold PBS with 5 mM EGTA (Carl Roth, Karlsruhe, Germany) with the help of a cell scraper. Tumor tissue was homogenized with a rotor-stator homogenizer (Ultra-Turrax T8, IKA-Werke, Staufen, Germany) in ice-cold PBS with 5 mM EGTA. Both, cell and tissue homogenates were centrifuged for ten minutes (4 °C, 200 × g), and supernatants were discarded. Pellets were resuspended in 5 mL of membrane isolation buffer (5 mM TrisHCl pH 7.6, 5 mM MgCl_2_, 1 mM EGTA, protease inhibitor cocktail cOmplete [Roche Applied Science, Penzberg, Germany]) with a glass dounce tissue grinder (Wheaton, Millville, USA) and further homogenized by moving the pestle up and down (approx. 30 times). After centrifugation for 30 minutes (4 °C und 40.000 g), the process was repeated, before cell and tissue homogenates were resuspended in 1 mL membrane isolation buffer, aliquoted and stored at −80 °C. For competitive binding, 5 µg of isolated membrane in 25 μL binding buffer (50 mM 4-(2-hydroxyethyl)-1-piperazineethanesulfonic acid (HEPES) pH 7.4, 5 mM MgCl^2^, 1 mM CaCl^2^, 0.5% BSA, protease inhibitor cocktail cOmplete) were incubated with increasing concentrations of non-radioactive peptide (2-fold concentrated in 50 µL) and 100,000 cpm of ^125^I-chemerin-9 (in 25 μL binding buffer). After one hour of incubation at 37 °C, the mixture was transferred to 96-well filter plates (MultiScreen^HTS^ FB Filter Plate 1.0 µm, Millipore, Billerica, USA), unbound peptide was withdrawn by suction and the plate was washed four times with cold washing buffer (50 mM Tris-HCl pH 7.4, 125 mM NaCl, 0.05% BSA). After drying the filter plate, 40 μL scintillation cocktail (Ultima Gold F, Perkin Elmer, Waltham, USA) per well was added and radioactivity was measured by liquid scintillation counting (MicroBeta^2^ Microplate Counter, PerkinElmer, Waltham, USA).

### Radiochemical labelling

Radiolabeling experiments were performed on a Modular Lab PharmTracer synthesis module (Eckert & Ziegler, Berlin, Germany) which allows fully automated cassette-based labeling of gallium tracers utilizing a pharmaceutical grade ^68^Ge/^68^Ga generator (GalliaPharm, 1.85 GBq, good manufacturing practice (GMP)-certified; Eckert & Ziegler GmbH, Berlin, Germany). Cassettes were GMP-certified and sterile. They were used without pre-conditioning of the cartridges. The gallium generator was eluted with aqueous HCl (0.1 M, 7 mL) and the eluate was purified on an ion-exchange cartridge followed by elution using 1 mL of 0.1 M HCl in acetone. The peptide-DOTA conjugate (50 µg from a stock solution 1 mg/mL in 10% DMSO, 90% water) was mixed with 500 µL 0.1 M HEPES buffer (pH 7) and heated for 500 s at 95 °C. After the reaction, the mixture was passed through a C-18 cartridge for purification and the tracer was eluted from the cartridge with ethanol. For injection, the resulting solution was diluted with saline to ≤ 10% ethanol.

### PET/MR imaging

Positron emission tomography (PET)/magnetic resonance imaging (MRI) (1 Tesla nanoScan PET/MRI, Mediso, Hungary) was performed at the Berlin Experimental Radionuclide Imaging Center (BERIC), Charité – Universitätsmedizin Berlin. A dedicated mouse whole-body coil was used for RF-transmission and signal receiving. Mice were anesthetized with general anaesthesia (1-2% isoflurane/0.5 L/min oxygen). Body temperature was maintained at 37 °C during the time of imaging by using a heated bed aperture. Anatomic whole-body MRI scans were acquired using a high-resolution T2-weighted 2D fast spin echo sequence (T2 FSE 2D) with the following parameters: TR = 8700 ms; TE = 103 ms; slice thickness/gap = 1.1 mm/ 0.1 mm; matrix = 256 × 256 mm; external averages = 5 and number of excitations = 2. PET scans in list mode were acquired either for 30 minutes one and two hours after intravenous injection or for 90 minutes of dynamic imaging starting directly before injection of 0.15 mL of tracer, corresponding to a ^68^Ga activity of approximately 20 MBq for static 1 h-images and 2 h-images and 15 MBq for kinetic imaging. To determine the effect of unlabeled ligand on the tumor uptake, 200 nmol CG34 peptide (50-fold excess) was co-injected. Static PET images were reconstructed from the raw data as one image (1 × 1800 s) and dynamic PET images were reconstructed with the image sequence 9 × 600 s. The uptake value (kBq/cm^3^) in the tumor tissue was determined by manual contouring of a volume of interest (VOI) of the PET images using PMOD 3.610 (PMOD Technologies, Zürich, Switzerland).

### Biodistribution studies

Tumor-bearing mice were injected with approximately 10 MBq of ^68^Ga-DOTA-peptide to the tail vein via a catheter. Mice were sacrificed and dissected one or two hours after injection. Tumor, blood, stomach, pancreas, small intestine, colon, liver, spleen, kidney, heart, lung, muscle and femur samples were weighed and uptake of radioactivity was measured by a gamma counter. To determine the effect of unlabeled ligand on the tumor uptake, 200 nmol CG34 peptide (100-fold excess) was co-injected.

### Statistical analysis

All statistical analyses were performed using GraphPad Prism 5.04. EC_50_/IC_50_ values were determined by nonlinear sigmoidal curve fitting with variable slope setting, and normalized response for competitive binding results. Multiple group comparisons were done by a two-way ANOVA and Bonferroni post hoc test. All presented data are based on independent experiments. Normalizations and statistics are further defined in each figure legend.

### Data availability

Numerical data for all experiments (xlsx file) have been deposited in an open data repository for public access: http://doi.org/10.5281/zenodo.2591417

## Results

### Characterization of an endogenously CMKLR1-expressing tumor mouse model

As models for tracer testing, the human breast cancer cell line DU4475, which endogenously expresses high levels of CMKLR, and the target-negative human lung cancer cell line A549 were chosen to induce subcutaneous xenograft tumors in nude mice. Analysis of *ex vivo* tumor tissue confirmed CMKLR1 expression in DU4475 xenografts by immunostaining, whereas A549 tumors showed no detectable antigen (Figure 1A). To confirm presence of receptor protein in these xenograft tissues with an independent method, radioligand binding studies were performed on membrane preparations of cultured cells and *ex vivo* tissue (Figure 1B). Receptor binding of ^125^I-labeled chemerin-9 in DU4475 cell membrane preparations (*in vitro*) led to values around 200 cpm (assessment of radioactivity by liquid scintillation counting) which could be significantly blocked (approx. 80 cpm) by an excess of the unlabeled peptide (1 µM). Membranes isolated from DU4475 tumors (ex vivo) bound less labeled peptide (approx. 115 cpm) but still exhibited specific binding, as 1 µM chemerin-9 clearly displaced about 50% of the bound activity (60 cpm). In contrast, membrane preparations from A549 cells xenograft tumors showed no specific binding.

**Figure 1:**
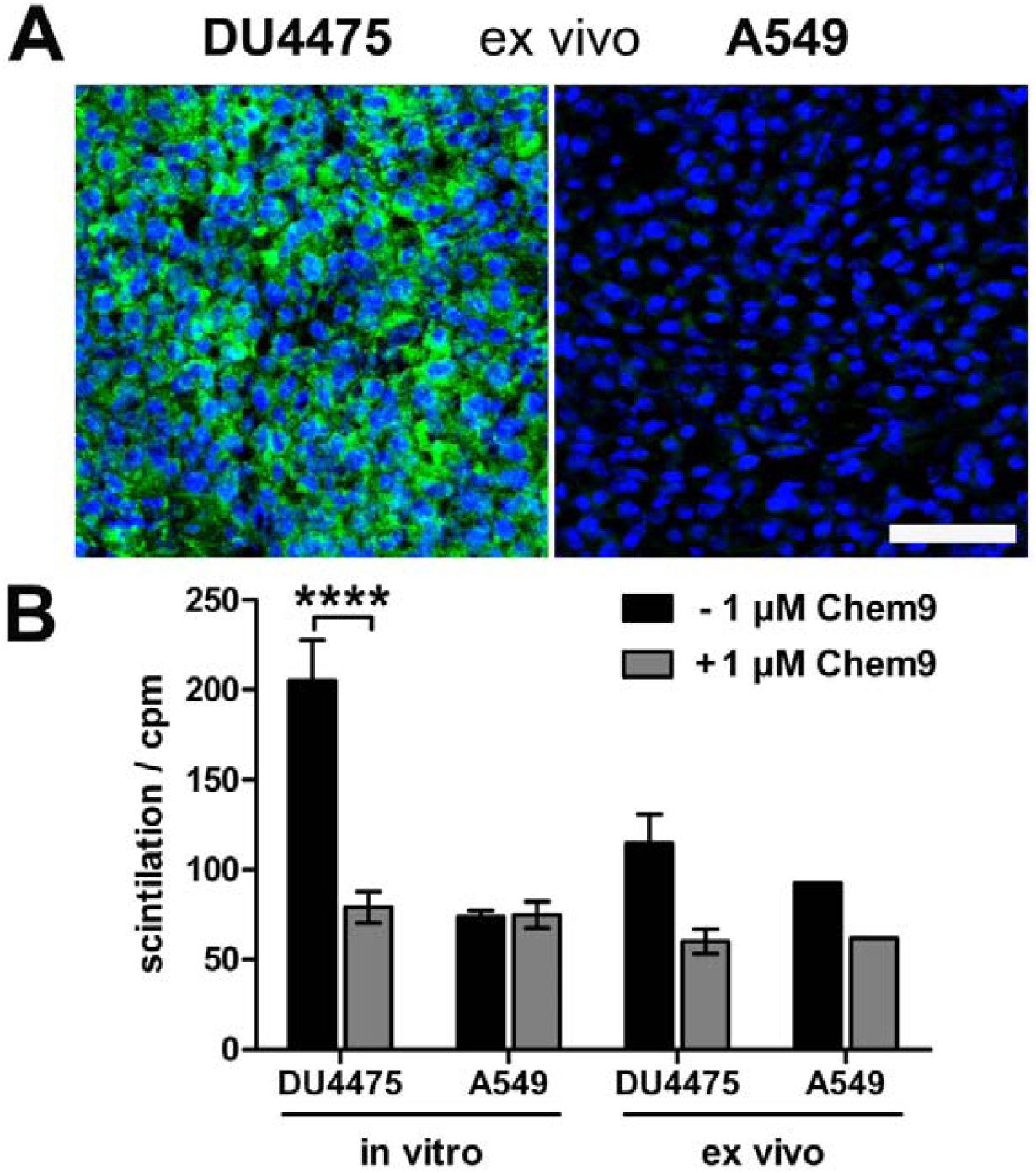
Characterization of the xenograft tumor model. Breast cancer DU4475 and lung cancer A549 cells were employed to establish a xenograft model with both CMKLR1-positive and -negative tumor. **(A)** Immunostaining of xenograft cryosections revealed CMKLR1 immunostaining (green) in DU4475 *ex vivo* tissue whereas the A549 tumor showed no signal. (blue: cell nuclei; bar = 50 µm) **(B)** Competitive radioligand binding studies with ^125^I-labeled chemerin-9 showed specific binding to DU4475 cell membranes (*in vitro*) and xenograft tumor membranes (*ex vivo*), which could be blocked by 1 µM unlabeled ligand (Chem9). In contrast, A549 cells showed no and A549 xenografts only minor binding. Data were obtained from at least three independent experiments and values are indicated as means ± SEM. (**** P < 0.0001)

### Characterization of high potency, high affinity CMKLR1-targeting peptide-DOTA conjugates

Chemical design of five tracers for CMKLR1 targeting involved two peptide analogs of the natural CMKLR1 ligand chemerin (CG34 or CG36) that were attached to the chelator 1,4,7,10-tetraazacyclododecane-1,4,7,10-tetraacetic acid (DOTA) via one of four different chemical linkers: 4,7,10-trioxatridecan-succinamic acid (TTDS), 6-aminohexanoic acid (AHX), N-ε-capryloyl-lysine (KCap) and 10-aminodecanoic acid (ADX) (Figure 2A). Due to their chemical composition, both peptide analogs as well as the four linkers exhibit distinct characteristics such as different length and hydrophobicity. In Figure 2A, these conjugate components are depicted in different colors (reddish colors indicate more hydrophobic properties and greenish colors more hydrophilicity).

**Figure 2:**
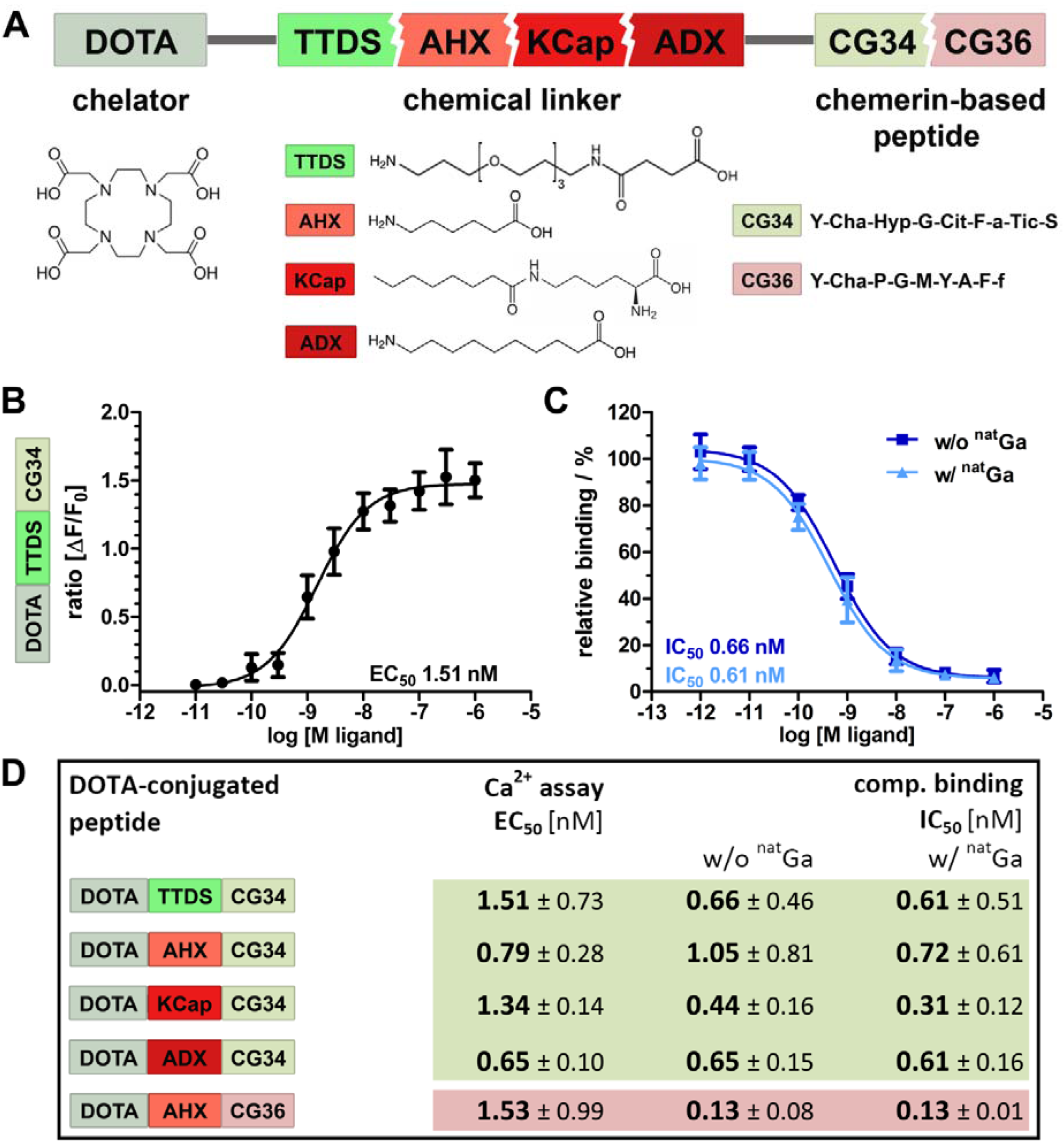
Characterization of DOTA peptide conjugates regarding their functionality and affinity. **(A)** The chelating agent 1,4,7,10-tetraazacyclododecane-1,4,7,10-tetraacetic acid (DOTA) was conjugated to the N-terminus of one of the two optimized chemerin peptide analogs, CG34 and CG36, by one of the four chemical linkers. **(B)** By functional Ca^2+^ mobilization assay, the concentration-response curve for the TTDS-linked CG34 conjugate could be determined and the resulting EC_50_ value (1.51 nM) confirmed its high receptor-activating potency. Response data are presented as mean ± SEM of three independent experiments. **(C)** Ligand affinity to the receptor was assessed by competitive radioligand binding experiments using ^125^I-chemerin-9 as radioligand. Displacement binding curves revealed similar IC_50_ values of the labeled (with ^nat^Ga) and unlabeled (without ^nat^Ga) DOTA conjugate indicating high affinity. Data are shown as mean ± SEM of three independent experiments normalized to non-competed radioligand binding. **(D)** To assess how the conjugation affects the ligand characteristics, functional and affinity data were obtained accordingly for all five chemerin conjugates (summarized as EC_50_/IC_50_ values). The Ca^2+^ and binding assays were performed with HEK293a cells stably expressing huCMKLR1 and G_α16_. All EC_50_/IC_50_ values were obtained from three independent experiments and are indicated as means ± SD. (green colors indicate more hydrophilic, red more hydrophobic properties of peptides and linker)

*In vitro* characterization of the probes was realized by activity and radioligand binding studies in HEK293a cells stably transfected with CMKLR1. Ligand-induced receptor activation as intracellular Ca^2+^ flux was quantified in an intracellular calcium mobilization assay (Figure 2B). Concentration-response profiles for all five tracers were determined and EC_50_ values were calculated. In comparison to the unconjugated peptide analog CG34 (EC^50^ 0.4 nM), the TTDS-linked DOTA conjugate of CG34 had a slightly higher EC_50_ value of 1.51 nM. Receptor affinity was assessed by competitive radioligand binding studies and resulting IC_50_ values were derived (Figure 2C). Furthermore, DOTA conjugates were labeled with non-radioactive gallium (^nat^Ga) to investigate a potential impact of metal ion complexation on binding affinity. For DOTA-TTDS-CG34, gallium labeling barely had an effect on receptor binding (IC_50_ value approximately 0.6 nM). As the unconjugated peptide CG34 exhibited an IC^50^ of 1.0 nM, conjugation with DOTA had even increased its binding affinity. Figure 2D summarizes all EC_50_ and IC_50_ values of the five DOTA peptide conjugates. All CG34 conjugates showed EC^50^ values within the low nanomolar range of approx. 0.6 – 1.5 nM, thus, 1.6-to 3.75-fold higher values for receptor activation than for the unconjugated peptide. However, for receptor binding the IC_50_ values were similar to the value for CG34 or even lower (subnanomolar range: approx. 0.3 – 1.0 nM) and cold gallium labeling did not adversely affect the affinity. Compared to the activating potency (EC_50_ 0.9 nM) and binding affinity (IC_50_ 0.1 nM) of the unconjugated peptide CG36, the DOTA-AHX-CG36 probe exhibited only slightly higher values (EC^50^ 1.53 nM, IC^50^ w/o ^nat^Ga 0.13 nM and w/ ^nat^Ga 0.13 nM).

### *In vivo* PET/MR imaging and biodistribution studies demonstrate specific tracer uptake in CMKLR1-expressing tumors

An initial PET study was performed to estimate the *in vivo* behavior of the five tracer conjugates after ^68^Ga labeling. Tumor-bearing mice received an intravenous injection of approximately 20 MBq radiotracer and were imaged for 30 minutes starting one hour post-injection in the PET/MRI scanner. Figure 3 shows *in vivo* PET images of the five tracers, depicted with hydrophobicity increasing from left to right. The most hydrophilic tracer ^68^Ga-DOTA-TTDS-CG34 accumulated mainly in the kidneys (Figure 3A, yellow arrow) with no clear DU4475 tumor uptake (right shoulder) compared to the general background. With ^68^Ga-DOTA-AHX-CG34, tumor-to-background signal was found to be higher (Figure 3B, white arrow) and accompanied by less kidney uptake. The tracer uptake within the target-positive tumor on the right shoulder was even more enhanced with KCap as linker (Figure 3C, white arrow). The most hydrophobic CG34 tracer, with ADX as a linker, also led to specific tumor uptake (Figure 3D, white arrow), but also high kidney (yellow arrow) and apparent liver uptake (red arrow). For the tracer ^68^Ga-DOTA-AHX-CG36, kidney and liver signals were high but almost no tumor uptake could be detected (Figure 3E). Based on these initial *in vivo* findings, the three most promising tracers ^68^Ga-DOTA-AHX-CG34, ^68^Ga-DOTA-KCap-CG34 and ^68^Ga-DOTA-ADX-CG34 were further analyzed. In addition to PET/MRI scans, *ex vivo* biodistribution studies were performed.

**Figure 3:**
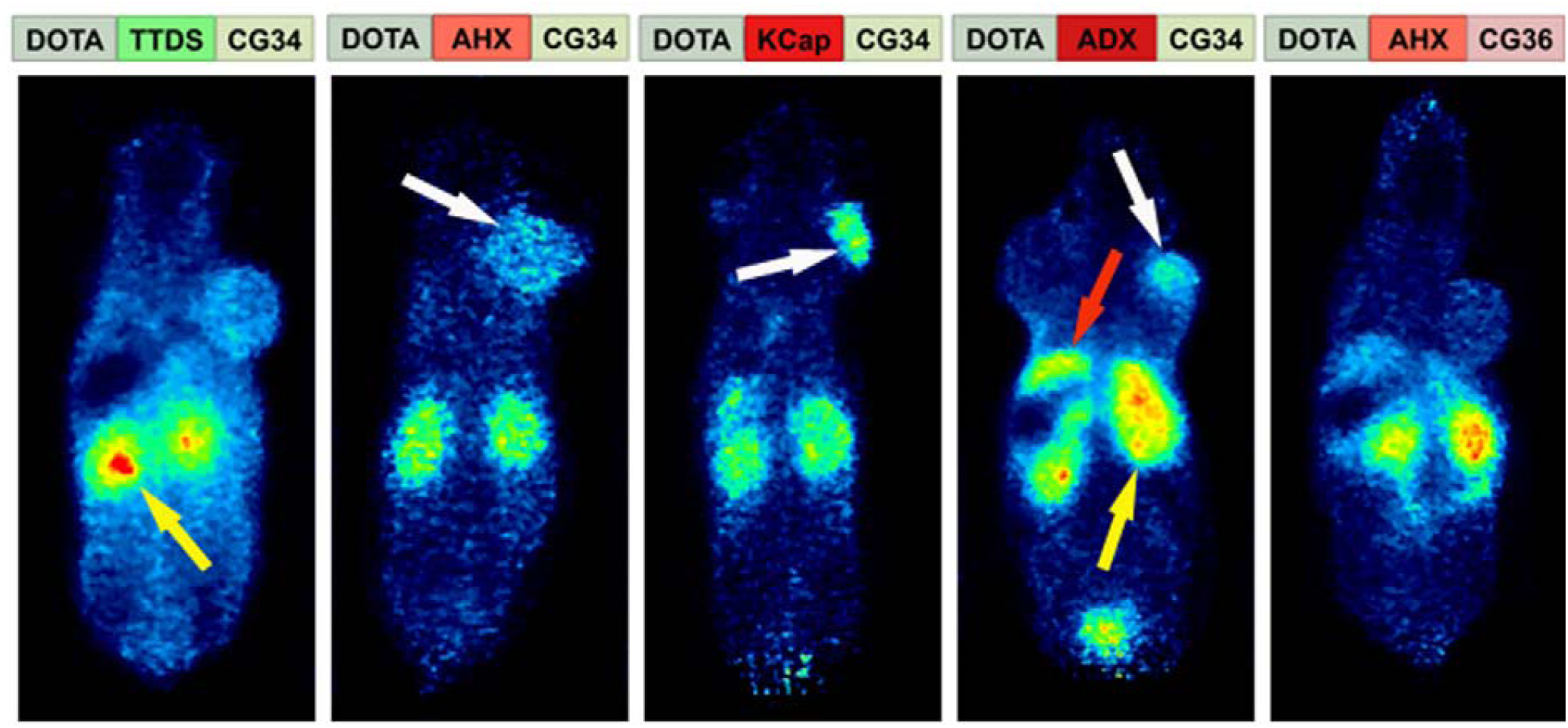
Comparison of tracer biodistribution by PET imaging. Representative static PET images were acquired for 30 minutes one hour post-injection of the five ^68^Ga-labeled DOTA conjugates in a DU4475/A549 tumor model and revealed a differential distribution. Approximately 20 MBq of each tracer were injected intravenously. **(A)** Predominant renal excretion of the hydrophilic tracer ^68^Ga-DOTA-TTDS-CG34 is indicated by a pronounced kidney signal (yellow arrow). **(B-C)** The more hydrophobic linkers AHX and KCap led to less kidney signal, but more distinct DU4475 uptake (white arrows). **(D)** Beside the kidney uptake, ^68^Ga-DOTA-ADX-CG34 induced an increased liver signal (red arrow). **(E)** ^68^Ga-DOTA-AHX-CG36 showed a comparable kidney and liver uptake, but less accumulation in the DU4475 tumor.

*In vivo* hybrid imaging with ^68^Ga-DOTA-KCap-CG34 and ^68^Ga-DOTA-ADX-CG34 using the PET/MRI scanner enabled not only functional PET analysis of the tracers but also allowed to gain anatomical insights into the tumor tissue and volume by high-resolution T2-weighted MR imaging (upper panels, Figure 4). As seen in the T2-weigthed images, both tumors (A549 on the left and DU4475 on the right shoulder) were of comparable size, tissue characteristics and vascularization. The KCap-linked radiotracer apparently accumulated in the DU4475 tumor and in kidneys one hour post-injection with decreasing PET signals after two hours. In contrast, no clear A549 tumor signal could be detected (upper panel, Figure 4A). In addition to *in vivo* imaging, tumor-bearing mice were injected with approximately 10 MBq of radiotracer and sacrificed after one and two hours, respectively. By *ex vivo* measurement, tissue radioactivity was calculated as percentage of injected activity per gram tissue (% IA/g). After one hour, ^68^Ga-DOTA-KCap-CG34 led to a DU4475 tumor uptake of about 4.5% IA/g and was 2-fold less in the A549 tumor with approx. 2.3% IA/g. Furthermore, higher values were measured for spleen (approx. 4.9% IA/g) and, due to the predominant renal excretion of peptides, for kidneys (approx. 5.8% IA/g). The DU4475 uptake decreased to approx. 2.9% IA/g, whereas the kidney signal increased up to approx. 6.5% IA/g two hours post-injection (lower panel, Figure 4A). For ^68^Ga-DOTA-ADX-CG34, the PET signal in DU4475 tumor and kidneys was comparatively stronger one and two hours post-injection than the uptake of ^68^Ga-DOTA-KCap-CG34 and there was also an additional distinct liver uptake of the radiotracer (upper panel, Figure 4B). These higher organ uptakes were confirmed by biodistribution data (lower panel, Figure 4B). In addition to overall higher values one hour after injection, liver (approx. 14.2% IA/g), kidney (10.3% IA/g) and spleen (7.8% IA/g) uptakes were the most profound off-target effects. However, with approx. 6.2% IA/g for CMKLR1-positive DU4475 tumors and approx. 2.7% IA/g for target-negative A549 tumors, PET results and tracer specificity could be confirmed. After two hours, DU4475 uptake declined by 1.5-fold to approx. 4.2% IA/g and a distinct liver uptake (approx. 12.5% IA/g) persisted. Kinetic measurements of tracer concentration in tumors, kidneys and liver demonstrated a delayed kidney peak for ^68^Ga-DOTA-KCap-CG34, the most hydrophobic molecule, with rapid washout from kidneys for both ^68^Ga-DOTA-AHX-CG34 and ^68^Ga-DOTA-ADX-CG34 (Figure S2). While the other two tracers showed a small decline in the DU4475 tumor, ^68^Ga-DOTA-ADX-CG34 appeared to gain in tumor activity until 90 min p.i.

**Figure 4:**
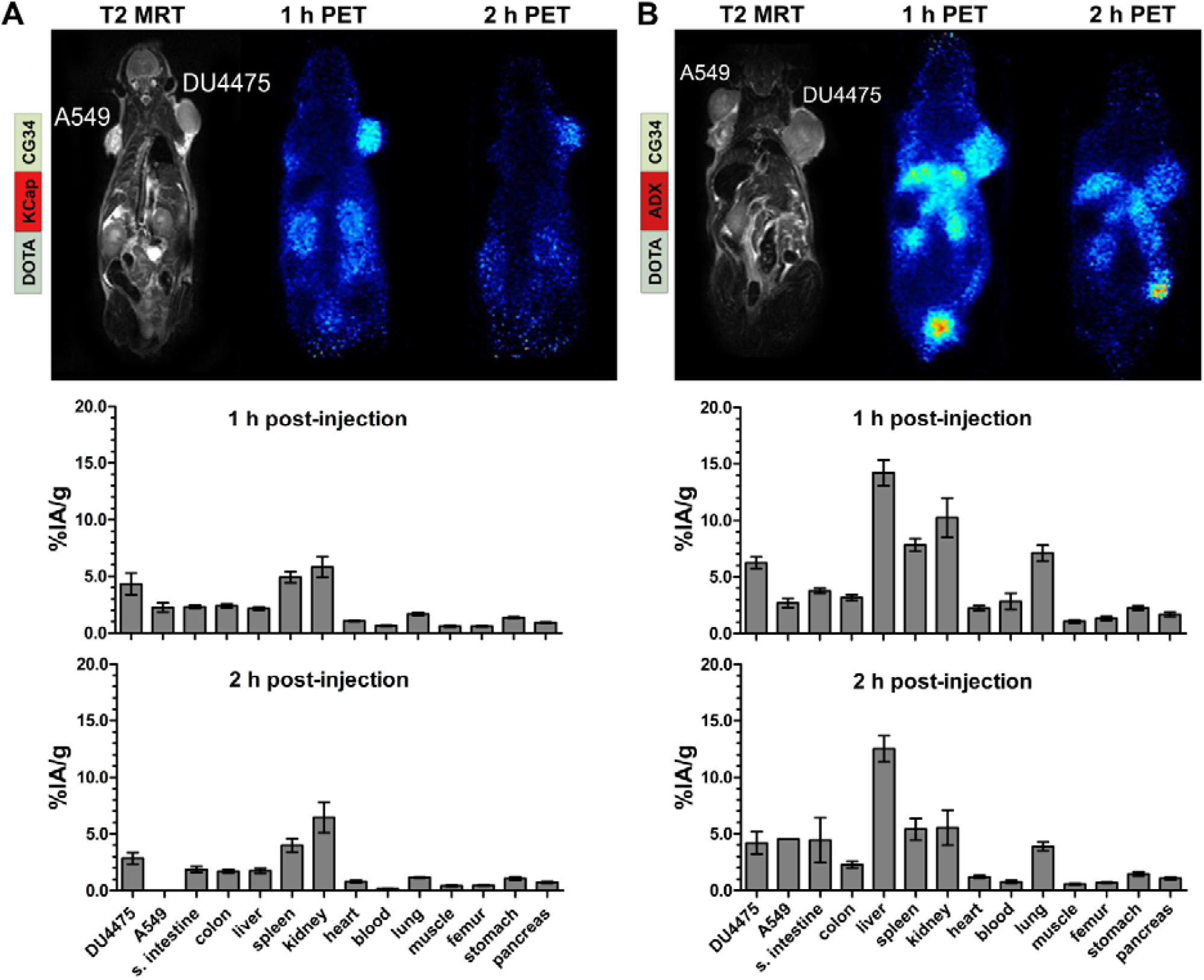
PET/MR imaging and biodistribution study with CMKLR1-targeted, chemerin-based tracers ^68^Ga-DOTA-KCap-CG34 and ^68^Ga-DOTA-ADX-CG34. Representative static PET images were acquired for 30 minutes one and two hours post-injection (p.i.) of 20 MBq ^68^Ga-labeled conjugate in DU4475 (target-positive) and A549 (negative) tumor model (upper panel). For quantitative analysis of tracer biodistribution, 10 MBq of the tracer was injected intravenously and tissue was analyzed *ex vivo* one or two hours p.i. Values for tracer uptake are indicated as percent injected activity per gram tissue (% IA/g) (lower panel). **(A)** The T2-weighted MRI image clearly shows both tumors on the animal’s shoulders. The PET images reveal ^68^Ga-DOTA-KCap-CG34 uptake in DU4475 after one hour, which decreased two hours p.i. The A549 tumor accumulated much less activity. Beside tumor uptake, the tracer accumulated in the kidneys after one hour. The quantitative analysis of biodistribution indicates the clear tracer uptake in DU4475 as compared to A549. Predominant renal excretion is represented by a noticeable kidney value (3 ≤ n ≤ 15; mean ± SEM). **(B)** Injection of ^68^Ga-DOTA-ADX-CG34 resulted in a strong DU4475 and low A549 uptake. In addition, tracer uptake was detected in kidneys and liver. Quantitative analysis of tracer biodistribution after one hour shows a high uptake in DU4475, and less in A549 tumors. Furthermore, the more hydrophobic tracer accumulated in the kidneys, liver and spleen and led to an overall stronger and longer organ accumulation (5 ≤ n ≤ 16; mean ± SEM).

### Receptor blocking studies verify tracer specificity

For further investigation of CMKLR1 specificity of the radiotracers, blocking experiments were carried out. To examine whether tracer binding to the tissues could be displaced, PET/MR imaging was performed twice with the same animal, once with and once without an excess of unlabeled peptide. In parallel, biodistribution experiments under blocking conditions (excess of unlabeled peptide) were performed to confirm PET results independently in a quantitative manner.

Figure 5 shows representative results of *in vivo* receptor blocking experiments with ^68^Ga-DOTA-AHX-CG34. PET/MR imaging without excess of the unconjugated and unlabeled peptide CG34 showed DU4475 tumor and kidney signals one hour post-injection (Figure 5A). However, co-injection of 200 nmol CG34 (50-fold excess) led to a complete loss of tumor uptake and a highly enhanced kidney signal (Figure 5B). The tracer biodistribution one hour after injection without competing CG34 clearly demonstrated the distinct DU4475 tumor uptake (approx. 5.1% IA/g) and the high kidney (approx. 6.5% IA/g) and spleen (approx. 6.3% IA/g) values (Figure 5C). CMKLR1 blocking with an excess of CG34 led to an overall low tissue and organ radioactivity except for strong kidney uptake (8.6% IA/g) and excretion into the urinary bladder. The tracer uptake within DU4475 tumors could be blocked by a factor of 6.4 to approximately 0.8% IA/g (Figure 5D).

**Figure 5:**
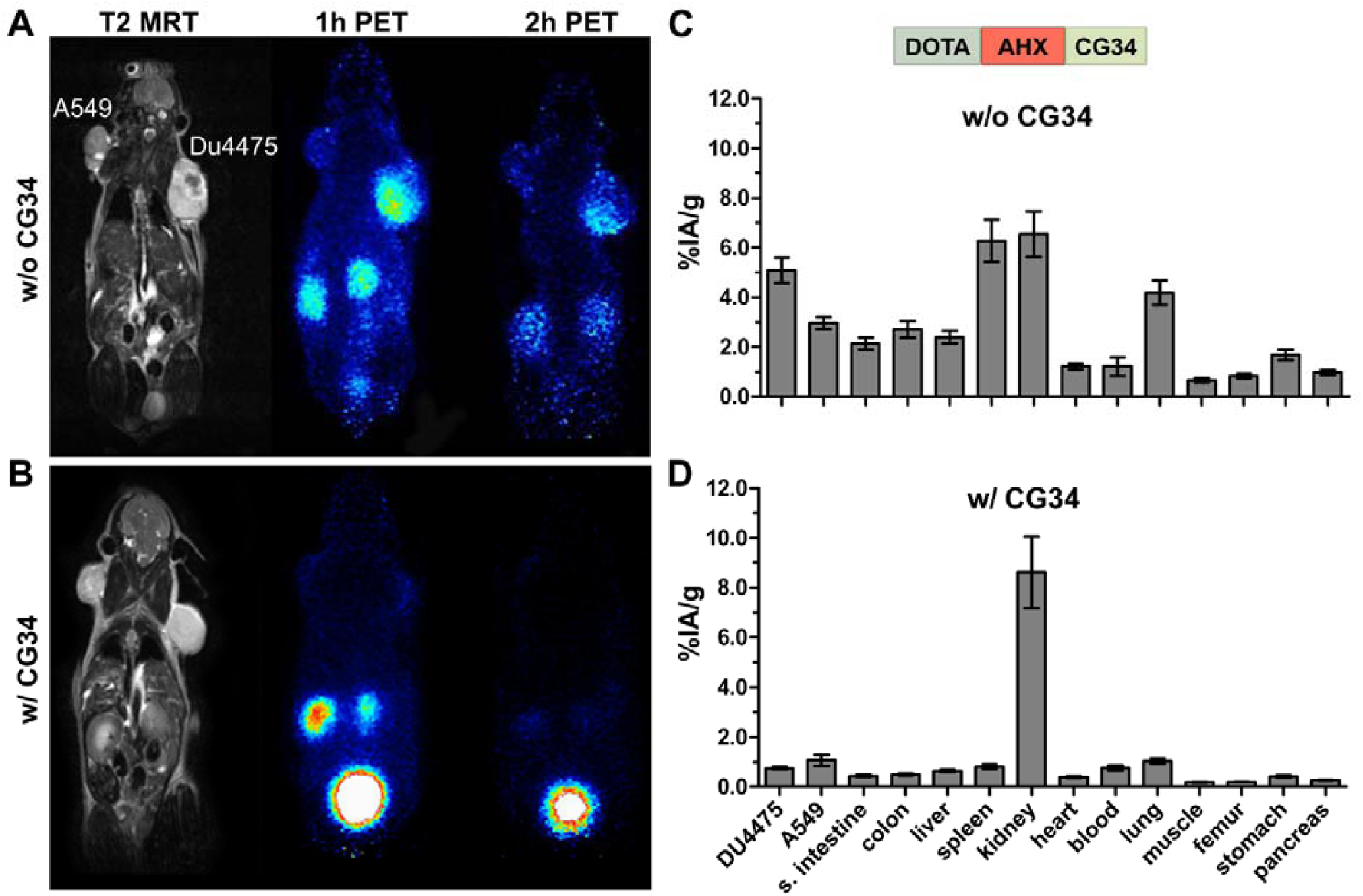
PET/MR imaging and biodistribution study along with a blocking experiment. Representative static PET images were acquired for 30 minutes one and two hours p.i. of ^68^Ga-labeled DOTA-AHX-CG34 conjugate **(A)** without and **(B)** with coinjected 50-fold excess of unlabeled peptide CG34 (200 nmol) in the same animal. 20 MBq of the tracer were injected intravenously. PET images show a strong DU4475 uptake, which could be completely displaced with excess of non-labeled peptide. Quantitative analysis of tracer biodistribution (10 MBq) one hour p.i. **(C)** without and **(D)** with receptor blocking confirmed mainly the specific tracer uptake in the DU4475 tumor. (6 ≤ n ≤ 13; mean ± SEM)

A summary of all biodistribution data including calculated tissue radioactivity ratios for the three most promising tracers ^68^Ga-DOTA-AHX-CG34, ^68^Ga-DOTA-KCap-CG34 and ^68^Ga-DOTA-ADX-CG34 is presented in Table 1. Use of these CMKLR1 tracers resulted in distinct target-specific DU4475 tumor uptake with 5.1% IA/g (^68^Ga-DOTA-AHX-CG34), 4.3% IA/g (^68^Ga-DOTA-KCap-CG34) and 6.2% IA/g (^68^Ga-DOTA-ADX-CG34). All three of them showed favorable tumor-to-kidneys ratios with 0.9, 0.7 and 0.8. CMKLR1 specificity could be proven by 1.8-fold (5.1 vs. 3.0% IA/g), 1.9-fold (4.3 vs. 2.3% IA/g) and 2.3-fold (6.2 vs. 2.7% IA/g) higher uptake in CMKLR1-positive DU4475 tumors than in target-negative A549 tumors one hour post-injection. In addition, receptor blocking by co-injection of an excess of unlabeled chemerin peptide diminished specific tracer binding by 6.4-fold (5.1 vs. 0.8% IA/g), 7.2-fold (4.3 vs. 0.6% IA/g) and 3.4-fold (6.2 vs. 1.8% IA/g). The most stable and durable tumor uptake was achieved by ^68^Ga-DOTA-AHX-CG34 with 4.9% IA/g (approx. 96% of the 1 h-value) after two hours, whereas uptake of the KCap-linked (approx. 67%) and ADX-linked (approx. 68%) tracers declined faster within two hours after injection. The highest overall non-tumor uptake values were seen with ^68^Ga-DOTA-ADX-CG34, with considerable liver (14.2% IA/g) and kidney (10.3% IA/g) uptake and hence, the lowest tissue radioactivity ratios for the DU4475 tumor.

**Table 1:** Biodistribution data and tissue radioactivity ratios of ^68^Ga-labeled chemerin tracers in the DU4475/A549 xenograft model. Data are presented as mean ± SEM (2 ≤ n ≤ 16) % IA/g of tissue or as a ratio. Blocking studies were performed in the presence of 200 nmol of CG34.

| Organ | <sup>68</sup>Ga-DOTA-AHX-CG34 |  |  | <sup>68</sup>Ga-DOTA-KCap-CG34 |  |  | <sup>68</sup>Ga-DOTA-ADX-CG34 |  |  |
| --- | --- | --- | --- | --- | --- | --- | --- | --- | --- |
|  | 1 h | 1 h-blocked | 2 h | 1 h | 1 h-blocked | 2 h | 1 h | 1 h-blocked | 2 h |
| DU4475 | 5.1 ± 0.5 | 0.8 ± 0.1 | 4.9 ± 0.8 | 4.3 ± 1.0 | 0.6 ± 0.1 | 2.9 ± 0.5 | 6.2 ± 0.5 | 1.8 ± 0.1 | 4.2 ± 1.0 |
| A549 | 3.0 ± 0.3 | 1.1 ± 0.2 | 1.4 ± 0.1 | 2.3 ± 0.4 | 0.6 ± 0.1 | n.d. | 2.7 ± 0.4 | 1.6 ± 0.2 | 4.5 |
| small intestine | 2.1 ± 0.2 | 0.4 ± 0.0 | 2.0 ± 0.4 | 2.3 ± 0.1 | 0.3 ± 0.1 | 1.9 ± 0.3 | 3.8 ± 0.2 | 1.4 ± 0.2 | 4.4 ± 2.0 |
| colon | 2.7 ± 0.3 | 0.5 ± 0.0 | 2.6 ± 0.7 | 2.4 ± 0.2 | 0.3 ± 0.0 | 1.7 ± 0.2 | 3.2 ± 0.3 | 1.2 ± 0.1 | 2.3 ± 0.3 |
| liver | 2.4 ± 0.3 | 0.6 ± 0.0 | 2.4 ± 0.4 | 2.2 ± 0.1 | 0.3 ± 0.0 | 1.8 ± 0.2 | 14.2 ± 1.1 | 10.4 ± 0.9 | 12.5 ± 1.1 |
| spleen | 6.3 ± 0.8 | 0.8 ± 0.1 | 5.0 ± 0.5 | 4.9 ± 0.5 | 0.5 ± 0.1 | 4.0 ± 0.6 | 7.8 ± 0.6 | 2.3 ± 0.4 | 5.4 ± 1.0 |
| kidney | 6.5 ± 0.9 | 8.6 ± 1.4 | 6.6 ± 1.1 | 5.8 ± 0.9 | 5.2 ± 0.9 | 6.5 ± 1.3 | 10.3 ± 1.7 | 14.5 ± 3.9 | 5.5 ± 1.6 |
| heart | 1.2 ± 0.1 | 0.4 ± 0.0 | 1.1 ± 0.3 | 1.1 ± 0.1 | 0.2 ± 0.0 | 0.8 ± 0.1 | 2.2 ± 0.2 | 1.4 ± 0.1 | 1.2 ± 0.1 |
| blood | 1.2 ± 0.4 | 0.8 ± 0.1 | 0.4 ± 0.1 | 0.6 ± 0.1 | 0.5 ± 0.1 | 0.2 ± 0.0 | 2.8 ± 0.7 | 3.2 ± 0.3 | 0.8 ± 0.1 |
| lung | 4.2 ± 0.5 | 1.0 ± 0.1 | 2.8 ± 0.3 | 1.7 ± 0.1 | 0.5 ± 0.1 | 1.2 ± 0.1 | 7.1 ± 0.7 | 3.4 ± 0.2 | 3.9 ± 0.4 |
| muscle | 0.7 ± 0.1 | 0.2 ± 0.0 | 0.5 ± 0.1 | 0.6 ± 0.1 | 0.2 ± 0.1 | 0.4 ± 0.1 | 1.1 ± 0.1 | 0.6 ± 0.1 | 0.6 ± 0.1 |
| femur | 0.8 ± 0.1 | 0.2 ± 0.0 | 0.7 ± 0.1 | 0.6 ± 0.1 | 0.2 ± 0.1 | 0.5 ± 0.1 | 1.3 ± 0.2 | 0.7 ± 0.1 | 0.7 ± 0.1 |
| stomach | 1.7 ± 0.2 | 0.4 ± 0.0 | 1.3 ± 0.1 | 1.4 ± 0.1 | 0.3 ± 0.1 | 1.1 ± 0.2 | 2.2 ± 0.2 | 1.1 ± 0.1 | 1.5 ± 0.2 |
| pancreas | 1.0 ± 0.1 | 0.3 ± 0.0 | 0.8 ± 0.1 | 0.9 ± 0.1 | 0.2 ± 0.0 | 0.7 ± 0.1 | 1.7 ± 0.2 | 0.8 ± 0.1 | 1.1 ± 0.1 |
| tumor-to-blood | 5.9 ± 0.7 | 1.0 ± 0.1 | 13.3 ± 3.0 | 7.3 ± 1.6 | 1.3 ± 0.3 | 18.2 ± 3.4 | 3.0 ± 0.3 | 0.6 ± 0.1 | 5.7 ± 1.1 |
| tumor-to-liver | 2.3 ± 0.2 | 1.2 ± 0.0 | 2.2 ± 0.5 | 2.1 ± 0.5 | 1.8 ± 0.3 | 1.8 ± 0.4 | 0.4 ± 0.0 | 0.2 ± 0.0 | 0.3 ± 0.1 |
| tumor-to-kidney | 0.9 ± 0.1 | 0.1 ± 0.0 | 0.7 ± 0.1 | 0.7 ± 0.1 | 0.1 ± 0.0 | 0.6 ± 0.1 | 0.8 ± 0.1 | 0.2 ± 0.0 | 0.8 ± 0.2 |
| tumor-to-pancreas | 5.6 ± 0.7 | 2.9 ± 0.2 | 6.2 ± 1.3 | 4.9 ± 1.2 | 3.7 ± 0.5 | 4.3 ± 0.9 | 3.9 ± 0.3 | 2.3 ± 0.2 | 3.9 ± 0.7 |
| tumor-to-muscle | 8.5 ± 1.0 | 4.4 ± 0.3 | 11.4 ± 2.4 | 8.9 ± 2.3 | 4.3 ± 0.9 | 7.2 ± 1.2 | 6.3 ± 0.5 | 3.3 ± 0.3 | 7.2 ± 1.2 |

## Discussion and Conclusions

The purpose of this study was to evaluate a family of five novel CMKLR1 tracers for their capacity to generate target-specific signals in a breast cancer xenograft mouse model. To the best of our knowledge, this is the first report of CMKLR1 *in vivo* imaging. Although known for its role in inflammation and metabolic regulation for many years, CMKLR1 and its chemokine-like ligand chemerin were only recently recognized as modulators of tumor proliferation and expansion. While other labs have reported overexpression and functional involvement of CMKLR1 in esophageal squamous cell carcinoma, hepatocellular carcinoma, neuroblastoma and gastric cancer [27-30], we have recently found CMKLR1 overexpression in breast cancer and pancreatic adenocarcinoma and have consequently developed two peptide analogs of chemerin for use in receptor-mediated tumor targeting (manuscript in preparation). These peptide analogs were based on chemerin-9, which has full agonistic activity towards the receptor CMKLR1 and corresponds to the C-terminal nine amino acids of processed chemerin [32]. In a previous study, we have shown the potential to generate stable and potent chemerin-9 analogs by the introduction of D-amino acids via an evolutionary algorithm [33]. The resulting peptide analogs showed improved properties concerning receptor activation, binding affinity and metabolic stability. In this study, we have made use of such analogs as components of chelator conjugates to do PET/MR imaging in a breast cancer xenograft mouse model.

The suitability of the animal model with CMKLR1-positive DU4475 and CMKLR1-negative A549 xenograft tumors was confirmed by immunofluorescence staining and radioligand binding studies. As expression of proteins in cells may change upon transfer from the culture dish monolayer to three-dimensional subcutaneous xenograft tumor tissue in mice, we investigated CMKLR1 expression in tumor tissue. Thereby, overexpression as well as ligand binding were established for DU4475 tumors while A549 tumors were confirmed as a negative control. The moderate tracer uptake seen in the latter is probably due to unspecific mechanisms, such as the enhanced retention and permeability effect or blood pool activity.

In the design of our tracers, we followed the hypothesis that tumor uptake and biodistribution may depend on a balance of hydrophobic and hydrophilic properties within the molecule. Therefore, we chose not only to make use of a more hydrophilic and a more hydrophobic CMKLR1 peptide ligand analog, but also to take advantage of different linkers to introduce a varying degree of hydrophobicity into the conjugates. As DOTA with its four carboxylic groups confers rather strong hydrophilic features, we chose to compensate this with mainly hydrophobic linker moieties: 6-aminohexanoic acid (AHX), N-ε-capryloyl-lysine (KCap) and 10-aminodecanoic acid (ADX). Only one linker displayed hydrophilic properties: the PEG-like 4,7,10-trioxatridecan-succinamic acid (TTDS) (Figure 2A). A set of five resulting DOTA peptide conjugates was tested *in vitro* prior to the animal study. Of note, all these tracers proved to be of high affinity and functional activity, with IC_50_ and EC_50_ values being below 2 nM (Figure 2D). Affinity also did not change significantly upon complexation of DOTA with non-radioactive gallium, and if so, IC_50_ values even decreased. This was in many ways unexpected as we and others had often experienced an impairment of the binding and activating capacity of ligands upon conjugation with a linker and a reporter such as a chelator or dye molecule [34-37]. Likewise, the affinity of such a conjugate may be affected by the coordination of a radiometal ion [38]. The structural requirements of chemerin analogs binding to CMKLR1 obviously allow for a broad variety of additions to the N-terminus of the nonameric peptide. Even though the two most potent conjugates were the ones with the shortest linkers (AHX and ADX), differences were small and a significant systematic influence of linker hydrophobicity or length on affinity or activity was not observed.

The initial PET studies using all five tracers revealed a clear difference regarding tumor uptake and biodistribution: the most hydrophilic conjugate (^68^Ga-DOTA-TTDS-CG34) and the most hydrophobic conjugate (^68^Ga-DOTA-AHX-CG36) both showed lower tumor uptake and higher kidney uptake than the three conjugates with balanced hydrophilicity in the molecule (Figure 3). This was also confirmed by results from biodistribution experiments: for both tracers, there was no clear difference in uptake between target-positive and target-negative tumor (Figure S1 and Table S1). These results may indicate the validity of our hypothesis that sufficient and specific tumor uptake may be achieved best using a conjugate with moderate, balanced hydrophilicity. Both highly hydrophobic and strongly hydrophilic tracers may primarily yield unspecific uptake in kidney and liver.

By peptide optimization and conjugation, we were able to obtain a family of five high-potency, high-affinity molecular probes for further *in vivo* application as radioactive tracers. In addition to PET/MRI scans one and two hours post-injection, *ex vivo* biodistribution studies were performed to assess quantitative values for tissue uptake. Utilizing a breast-cancer xenograft mouse model, PET/MRI and concomitant biodistribution studies revealed the specificity of our chemerin-based radiotracers by distinct PET signals and uptake values of CMKLR1-positive DU4475 tumors compared to no PET visibility and low tumor radioactivity values of target-negative A549 tumors.

Kidney uptake often limits therapeutic use of tracers. For the three best tracers described in this study, renal uptake was between 5.8 to 10.3% IA/g, resulting in favorable tumor-to-kidney ratios of 0.7 to 0.9 at one hour post injection. Further evidence for the specificity of the tracer was obtained in a blocking experiment with an excess of non-labeled peptide (Figure 5). As obvious from both PET/MR images as well as biodistribution data, this excess strongly reduced uptake of the tracer ^68^Ga-DOTA-AHX-CG34 in CMKLR1-positive DU4475 tumors, indicating displacement of the radioligand from its receptor. The lower uptake observed in other organs probably also corresponds to tracer displacement from chemerin receptors known to be expressed there (e.g. lung, spleen) [21, 39]. The dynamic PET scan of ^68^Ga-DOTA-AHX-CG34 showed faster background/off-target clearance as well as fast and stable accumulation at tumor sites. Whereas higher hydrophobicity of KCap and ADX led to delayed and prolonged kidney uptake, possibly due to plasma protein binding or metabolization in the liver (Figure S2).

While a tumor uptake of 4-6% represents a promising targeting result for these first CMKLR1 tracers, several options for an improvement remain. One current limitation, especially for an application in peptide receptor radionuclide therapy (PRRT), is the fast elimination of the signal from the target-positive tumor. According to biodistribution data, the signal decrease between one hour and two hours in DU4475 tumors was about 25-70%. Although the underlying peptide analogs have been selected for their improved stability in *in vitro* protease degradation assays, a degradation due to proteolytic activity in circulation and target tissue may play a role. Similarly, fast excretion via the renal pathway, as well as hepatic degradation (e.g. ^68^Ga-DOTA-ADX-CG34) may create unfavorable distribution kinetics. The precise causes for this rapid decrease in tumor signal will have to be determined in subsequent studies. Biochemical isolation and analysis of tracer metabolites from the animal’s circulation may provide a means of clarifying the underlying mechanism.

## Abbreviations

% IA/g: percent injected activity per gram tissue
^18^F-FDG: ^18^F-fluoro-deoxyglucose
ADX: 10-aminodecanoic acid
AHX: 6-aminohexanoic acid
BERIC: Berlin Experimental Radionuclide Imaging Center
BSA: bovine serum albumin
CG34: chemerin analog with amino acid sequence Y-Cha-Hyp-G-Cit-F-a-Tic-S
CG36: chemerin analog with amino acid sequence Y-Cha-P-G-M-Y-A-F-f
Chem9: chemerin-9
CMKLR1: chemokine-like receptor 1
DMSO: dimethyl sulfoxide
DOTA: 1,4,7,10-tetraazacyclododecane-1,4,7,10-tetraacetic acid
DTPA: diethylenetriaminepentaacetic acid
EC^50^: half maximal effective concentration
ECM: extracellular matrix
EGFR: epithelial growth factor receptor
EGTA: ethylene glycol-bis(β-aminoethyl ether)-N,N,N′,N′-tetraacetic acid
ESCC: esophageal squamous cell carcinoma
G418: geneticin
GMP: good manufacturing practice
GPCR: G protein-coupled receptor
HEPES: 4-(2-hydroxyethyl)-1-piperazineethanesulfonic acid
HPLC: high-performance liquid chromatography
IC^50^: half maximal inhibitory concentration
KCap: N-ε-capryloyl-lysine
MRI: magnetic resonance imaging
NET: neuroendocrine tumor
NMRI: Naval Medical Research Institute
PBS: phosphate-buffered saline
PEG: polyethylene glycol
PET: positron emission tomography
PRRT: peptide receptor radionuclide therapy
PSMA: prostate-specific membrane antigen
RPMI1640: Roswell Park Memorial Institute medium 1640
SD: standard deviation
SEM: standard error of the mean
SSA: somatostatin analog
SSTR: somatostatin receptor
T2 FSE 2D: high-resolution T2-weighted 2D fast spin echo sequence
TBS: tris-buffered saline
TE: echo time
TFA: trifluoroacetic acid
TR: repetition time
TTDS: 4,7,10-trioxatridecan-succinamic acid
VOI: volume of interest

## Author Contributions

Conception and design: S.E., C.G. Development of methodology: S.E., C.G. Acquisition of data: S.E., L.N., E.J.K., J.D.C.G., S.P., A.W., J.L.v.H., S.H., S.E., S.B. Analysis and interpretation of data: S.E., C.G. Writing, review, and/or revision of the manuscript: S.E., E.J.K., N.B., W.B., C.G. Study supervision: W.Br., C.G.

