## Supplements for "CMKLR1-targeting peptide tracers for PET/MR imaging of breast cancer"

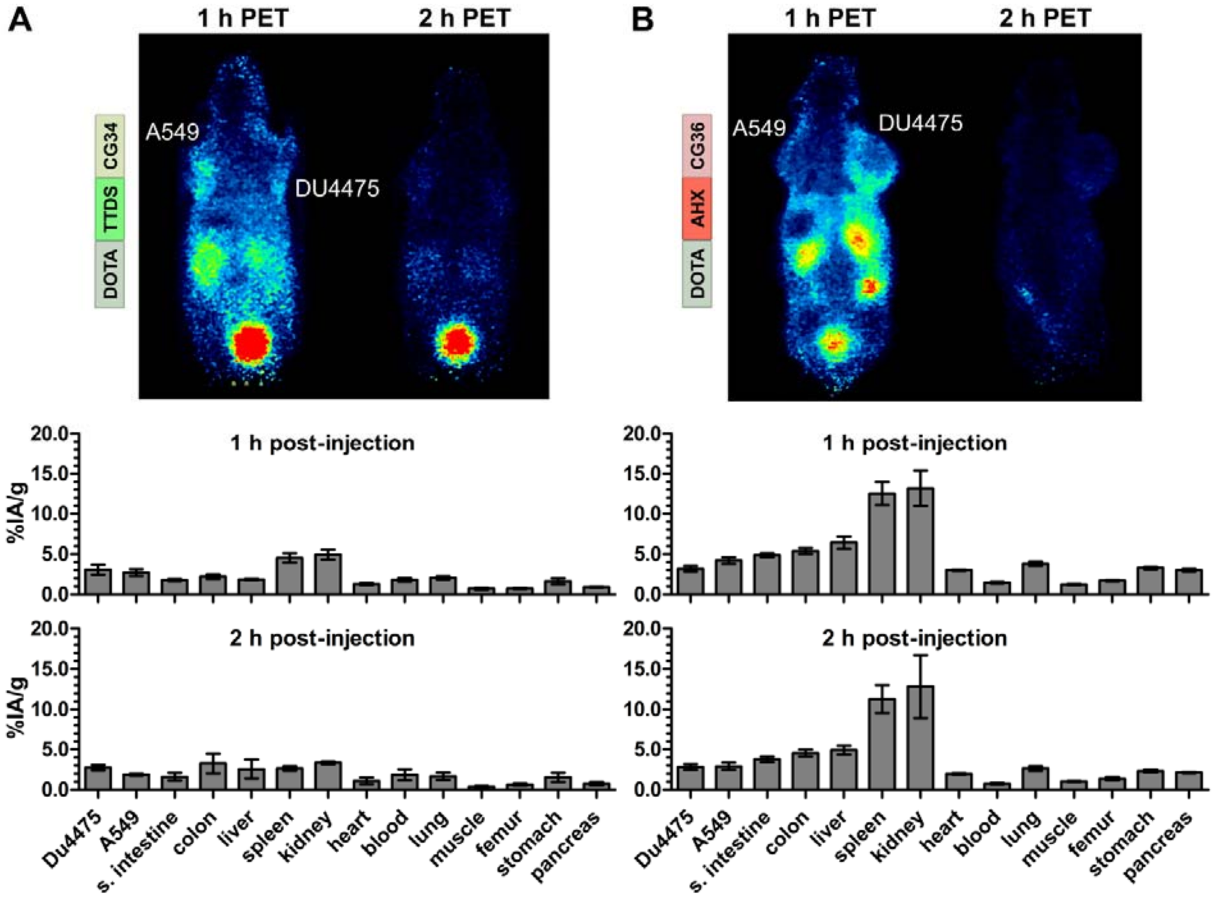


**Figure S1: PET/MR imaging and biodistribution study with CMKLR1-targeted, chemerin-based tracer $^{68}$Ga-DOTA-TTDS-CG34 and $^{68}$Ga-DOTA-AHX-CG36.** Representative, static PET images were acquired for 30 minutes one and two hours post-injection (p.i.) of 20 MBq $^{68}$Ga-labeled conjugate in DU4475 (target-positive) and A549 (negative) xenograft tumor model (upper panel). For quantitative analysis of tracer biodistribution, 10 MBq of the tracer were injected intravenously and tissue was analyzed *ex vivo* one or two hours p.i. Values for tracer uptake are indicated as percent injected activity per gram tissue (% IA/g) (lower panel). **(A)** Representative PET images one hour p.i. show no clear $^{68}$Ga-DOTA-TTDS-CG34 uptake in neither A549, nor DU4475 tumor as well as a low kidney accumulation. After two hours, barely any signal is left. Quantitative analysis of biodistribution confirms the overall low tissue radioactivity ($2 \leq n \leq 10$; mean ± SEM). **(B)** Injection of $^{68}$Ga-DOTA-AHX-CG36 resulted in stronger signals within the DU4475 tumor, the kidneys and the liver. Quantitative analysis of tracer biodistribution shows an overall higher and prolonged tissue radioactivity after one and two hours. This more hydrophobic tracer mostly accumulated in spleen and kidneys, but with no apparent DU4475 uptake ($2 \leq n \leq 8$; mean ± SEM).

**Table S1: Biodistribution data and tissue radioactivity ratios of $^{68}$Ga-labeled chemerin tracer DOTA-TTDS-CG34 and DOTA-AHX-CG36 in DU4475/A549 xenograft model.** Data are presented as mean ± SEM (2 ≤ n ≤ 10) % IA/g of tissue; blocking studies were performed in the presence of 200 nmol of CG34.

| Organ | $^{68}$Ga-DOTA-TTDS-CG34 | | | $^{68}$Ga-DOTA-AHX-CG36 | | |
|---|---|---|---|---|---|---|
| | 1 h | 1 h-blocked | 2 h | 1 h | 1 h-blocked | 2 h |
| DU4475 | 3.1 ± 0.6 | 0.7 ± 0.1 | 2.7 ± 0.3 | 3.2 ± 0.3 | 1.3 ± 0.1 | 2.8 ± 0.4 |
| A549 | 2.7 ± 0.4 | 1.1 ± 0.4 | 1.9 ± 0.1 | 4.2 ± 0.4 | 1.2 ± 0.2 | 2.9 ± 0.5 |
| small intestine | 1.8 ± 0.1 | 0.4 ± 0.1 | 1.6 ± 0.5 | 4.8 ± 0.2 | 1.8 ± 0.3 | 3.8 ± 0.4 |
| colon | 2.2 ± 0.3 | 0.5 ± 0.1 | 3.2 ± 1.2 | 5.4 ± 0.4 | 1.4 ± 0.3 | 4.5 ± 0.4 |
| liver | 1.8 ± 0.1 | 0.4 ± 0.0 | 2.6 ± 1.2 | 6.4 ± 0.8 | 2.2 ± 0.3 | 4.9 ± 0.6 |
| spleen | 4.5 ± 0.6 | 0.5 ± 0.1 | 2.6 ± 0.3 | 12.5 ± 1.4 | 2.0 ± 0.6 | 11.3 ± 1.7 |
| kidney | 4.9 ± 0.6 | 4.8 ± 0.9 | 3.4 ± 0.2 | 13.2 ± 2.2 | 10.7 ± 3.2 | 12.8 ± 3.9 |
| heart | 1.3 ± 0.1 | 0.5 ± 0.1 | 1.1 ± 0.4 | 3.0 ± 0.1 | 1.0 ± 0.1 | 2.0 ± 0.1 |
| blood | 1.8 ± 0.2 | 1.3 ± 0.2 | 1.9 ± 0.7 | 1.5 ± 0.1 | 1.3 ± 0.3 | 0.7 ± 0.1 |
| lung | 2.0 ± 0.2 | 0.7 ± 0.2 | 1.7 ± 0.4 | 3.8 ± 0.3 | 1.7 ± 0.2 | 2.6 ± 0.3 |
| muscle | 0.7 ± 0.1 | 0.2 ± 0.1 | 0.4 ± 0.1 | 1.2 ± 0.1 | 0.8 ± 0.2 | 1.0 ± 0.1 |
| femur | 0.7 ± 0.1 | 0.3 ± 0.1 | 0.6 ± 0.1 | 1.7 ± 0.1 | 1.0 ± 0.5 | 1.4 ± 0.2 |
| stomach | 1.6 ± 0.4 | 0.4 ± 0.1 | 1.5 ± 0.6 | 3.3 ± 0.2 | 1.0 ± 0.2 | 2.3 ± 0.2 |
| pancreas | 0.9 ± 0.1 | 0.3 ± 0.1 | 0.7 ± 0.2 | 3.0 ± 0.2 | 1.2 ± 0.2 | 2.1 ± 0.1 |
| tumor-to-blood | 2.0 ± 0.4 | 0.6 ± 0.2 | 2.7 ± 1.3 | 2.3 ± 0.3 | 1.1 ± 0.2 | 3.9 ± 0.6 |
| tumor-to-liver | 1.6 ± 0.3 | 1.9 ± 0.4 | 2.0 ± 0.9 | 0.5 ± 0.1 | 0.6 ± 0.0 | 0.6 ± 0.1 |
| tumor-to-kidney | 0.7 ± 0.2 | 0.2 ± 0.0 | 0.8 ± 0.1 | 0.3 ± 0.1 | 0.2 ± 0.0 | 0.3 ± 0.1 |
| tumor-to-pancreas | 3.2 ± 0.5 | 3.0 ± 0.9 | 5.1 ± 1.9 | 1.1 ± 0.1 | 1.2 ± 0.2 | 1.3 ± 0.2 |
| tumor-to-muscle | 4.5 ± 0.9 | 4.4 ± 1.5 | 8.9 ± 2.9 | 2.6 ± 0.3 | 1.9 ± 0.4 | 2.7 ± 0.3 |

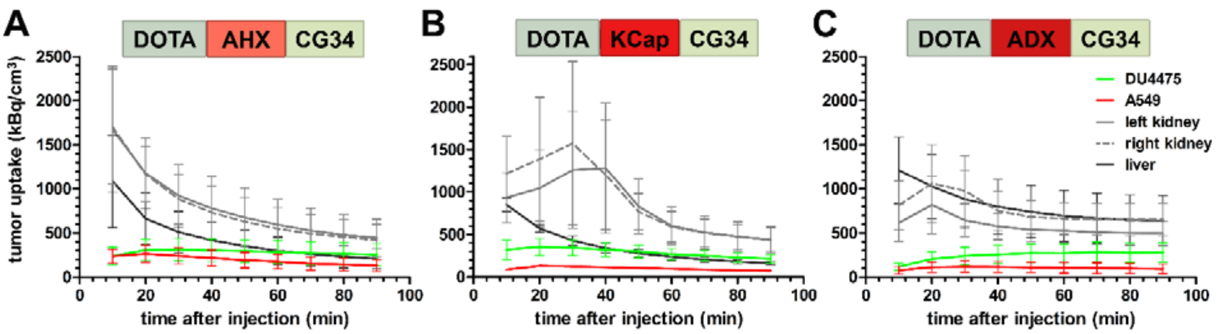


**Figure S2: PET signal kinetics of $^{68}$Ga-labeled CMKLR1 tracers in DU4475/A549 xenograft model.** Quantitative VOI analysis of tumor tissue, kidneys and liver depicting activity concentration kinetics derived from dynamic PET scanning. Two to three animals were injected with equal amounts of either tracer (15 MBq) immediately after starting the PET scan. Mean activity concentration [kBq/cm$^3$] from VOI analysis is shown as mean ± SEM.
